# Real-time fMRI-triggered experience-sampling: a proof-of-concept study

**DOI:** 10.64898/2026.05.08.723780

**Authors:** Tiara Bounyarith, David Braun, Aaron Kucyi

**Author notes:** **Correspondence to:** Aaron Kucyi, Ph.D. Dept. of Psychological & Brain Sciences, Drexel University 3201 Chestnut St. Room 304 Philadelphia, PA 19104 United States.

## Abstract

Much of a typical individual’s mental life is characterized by spontaneous thoughts that occur independently of external stimuli. In prior studies, ongoing mental experiences and their neural correlates have been captured using thought probes presented at random intervals during functional Magnetic Resonance Imaging (fMRI). However, this approach results in temporally imprecise estimates of brain activity relative to the arising of mental experience. In this preregistered, proof-of-concept study, we aimed to improve temporal precision using a novel method termed real-time fMRI-triggered experience-sampling (rt-fMRI-ES). We analyzed blood-oxygenation-level-dependent signals in real time during a wakeful resting state (n=60) to trigger thought probes from spontaneous activations within two regions: the dorsal anterior insular cortex (daIC; a key region within salience network) and posteromedial cortex (PMC; a key region within default mode network). We tested two preregistered hypotheses: (*<u>H1</u>*) Ratings of arousal time-locked to daIC-activation trials are higher than ratings time-locked to non-daIC-activation trials; (*<u>H2</u>*) Ratings of external-attention time-locked to PMC-activation trials are lower than ratings time-locked to non-PMC-activation trials. After applying preregistered exclusion criteria, 42 participants (1243 trials) and 49 participants (1429 trials) were included in H1 and H2 analyses, respectively. We did not find evidence in support of *H1*, but we did find evidence in support of *H2*, as external-attention ratings were significantly lower for trials triggered by PMC activation compared to other trial types. Taken together, we successfully developed and validated the rt-fMRI-ES method, offering a novel technique to efficiently capture spontaneous thoughts based on ongoing neural activity.

**Preregistered Stage 1 Recommendation:** https://osf.io/sd4hu (Date of in-principle acceptance: 07/24/2024; under temporary private embargo)

## 1 Introduction

Much of a typical individual’s mental life is characterized by involuntary thoughts, such as mind wandering, that occur independently of external stimuli (Kane et al., 2007, 2017; Killingsworth & Gilbert, 2010; Watkins & Roberts, 2020). There is growing scientific and clinical interest in understanding how the brain spontaneously generates such unprompted, inner mental experiences (Christoff et al., 2016; Kam et al., 2022; Kucyi et al., 2023; Smallwood et al., 2021). Functional neuroimaging studies have established a key role of the default mode network (DMN), and its interactions with other large-scale networks, in mind-wandering, spontaneous thought, and rumination (Andrews-Hanna et al., 2014; Berman et al., 2011; Mittner et al., 2016). Recently, involuntary thoughts have been measured during fMRI scans using online experience-sampling, where participants repeatedly provide ratings of their internal mental experiences at random time points throughout a scan. This method has yielded insights into the contents and dynamics of a wide array of ongoing, stimulus-independent experiences and their neural correlates (Christoff et al., 2009; Kucyi et al., 2013, 2021; Sormaz et al., 2018; Stawarczyk et al., 2011). However, it remains unknown how precisely ongoing mental events map on to distinct time points in spontaneous brain activity (Gonzalez-Castillo et al., 2021; Kucyi, 2018).

One of the major limitations of neuroimaging studies using the online experience-sampling method is the delivery of thought probes at random time intervals. This approach only captures a sliver of the abundance of unprompted mental experiences that an individual is bound to have during a resting state or task performance conditions. As random-interval experience-sampling is non-targeted in its execution, it has not aided in the investigation of when exactly a thought trajectory begins and ends. Furthermore, it can often miss the occurrence of specific thought subtypes, such as high-arousal, distressing mental experiences, that may be of interest in the context of mental health. Experience-sampling at random intervals leads to high variability in the temporal stages of sampled mental experiences. This often becomes a source of noise that necessitates large sample sizes in order to detect brain-experience relationships with typical group-averaging procedures (Kucyi et al., 2024). An alternative “self-caught” experience-sampling method has been tested during fMRI wherein individuals report the spontaneous arising of a thought (Ellamil et al., 2016; Hasenkamp et al., 2012). However, that approach only captures thought onsets that individuals are aware of and does not guarantee a uniform delay across trials or individuals between self-report and the brain activity that represents thought onset.

Here, to address these critical limitations, we develop and validate **real-time fMRI-triggered experience-sampling** (rt-fMRI-ES), a method for efficiently sampling self-generated thoughts based on spontaneous neural events. Using real-time analysis of incoming fMRI data during acquisition, we detect instances of high blood-oxygenation-level-dependent (BOLD) activation to trigger the appearance of thought probes immediately following neural events of interest. This approach aims to match the temporal stage of arising thoughts across trials and to efficiently target thoughts that have specific qualities, such as distressing experiences that underlie maladaptive disordered thought. Moreover, by treating brain activity as an independent variable, our approach aims to empower the prospective testing of brain-experience relationships. As the current study aimed to validate this novel rt-fMRI-ES approach, we additionally included trials that were not triggered by brain activity but were displayed after ample time had passed since the previous trial (termed “time-out trials”). This was to ensure that, in the event that an insufficient number of trials were triggered by brain activity in a given run, thought ratings were still being collected from participants.

We aimed to develop and validate rt-fMRI-ES by targeting both the **dorsal anterior insular cortex** (daIC), a core region of the brain’s salience (or cingulo-opercular) network (Seeley et al., 2007; Uddin et al., 2023), and the **posteromedial cortex** (PMC), a component of the DMN (Buckner & DiNicola, 2019; Raichle et al., 2001). The daIC is hypothesized to assign salience to self-generated experiences (Christoff et al., 2016), and its spontaneous activation has shown to be co-occur with neurophysiological markers of arousal (Kucyi & Parvizi, 2020; Schneider et al., 2016). Coupled with its atypical structure, function, and connectivity in psychiatric illness at a transdiagnostic level (Goodkind et al., 2015; Sha et al., 2019), there is a need to understand the role the daIC plays in characterizing distressing self-generated thought. In line with evidence of the daIC’s role in salient experiences, we predicted that subjective ratings of arousal time-locked to daIC-activation trials would be higher than ratings time-locked to non-daIC-activation trials (including both PMC-activation trials and time-out trials) *(H1)*. The PMC, which includes subregions of posterior cingulate cortex and precuneus, activates during internal mentation and self-related mental representations, and deactivates during explicit goal-directed attention states (Buckner & DiNicola, 2019; Cavanna & Trimble, 2006). Given the PMC’s role in self-referential and stimulus-independent thought, regardless of specific content, we predicted that subjective ratings of external-attention time-locked to PMC-activation trials would be lower than ratings time-locked to non-PMC-activation trials (including both daIC-activation trials and time-out trials) *(H2*) (see **Figure 1C**).

**Figure 1.**
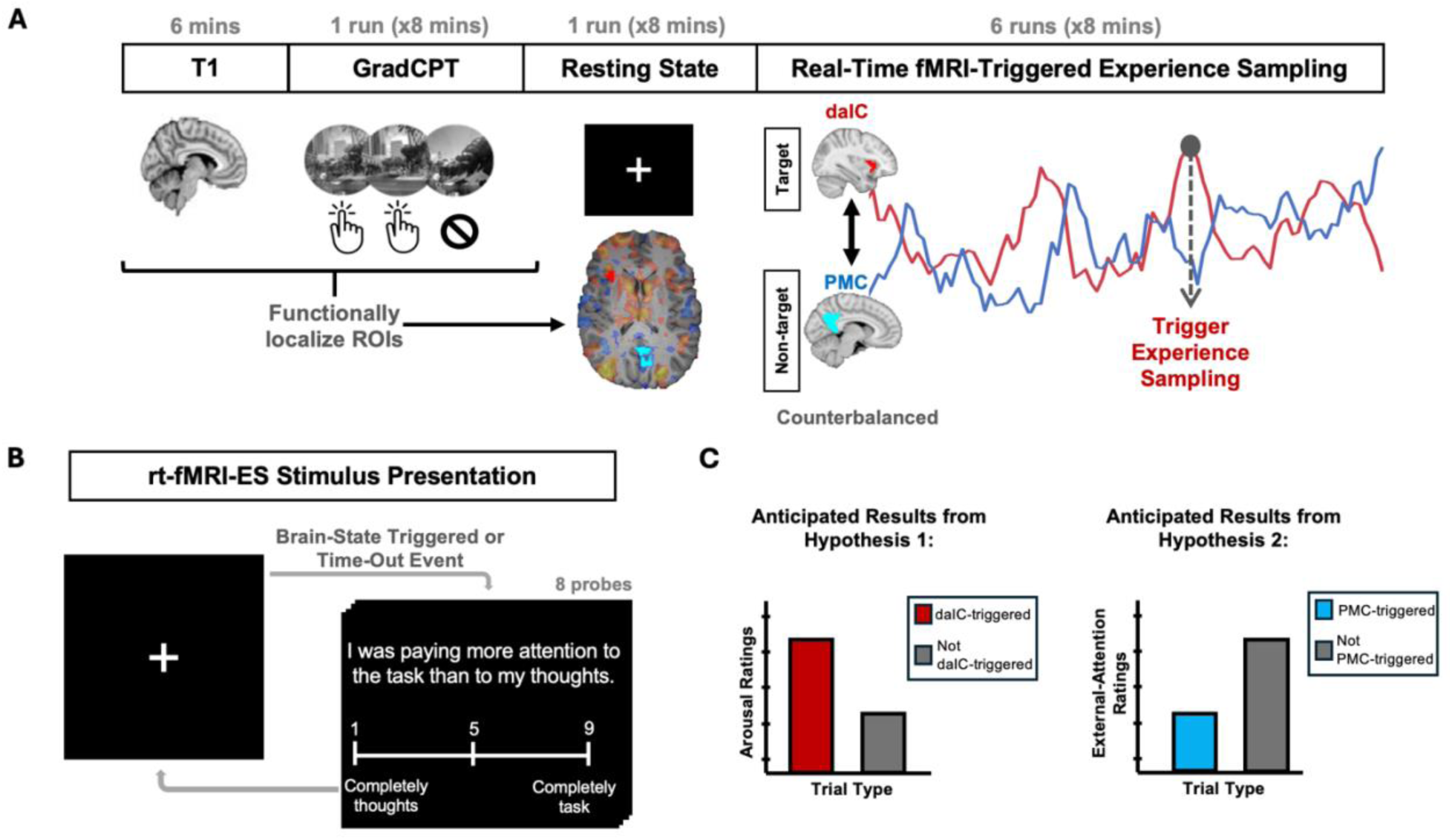
Overview of study procedures and task. **A)** Within a single neuroimaging session, participants underwent a T1 structural scan followed by an fMRI run while performing the Gradual-onset Continuous Performance Task (GradCPT). Experimenters used stimulus timing information from the GradCPT in conjunction with the participant’s structural and task fMRI images to compute individually localized regions of interest (ROIs) while the participant underwent a resting-state fMRI run. During GradCPT infrequent mountain events, the sample activation map produced by the functional localizer depicts whole-brain activation in transparent red-yellow and whole-brain deactivation in transparent blue. Individually-localized ROIs are shown in bright red (daIC) and bright light blue (PMC). Subsequently, across six runs of a real-time fMRI condition, brain activity within these individualized functional ROIs—the dorsal anterior insular cortex (daIC) and the posteromedial cortex (PMC)—was analyzed in real-time, with ROIs being counter-balanced across runs. Experience sampling trials were triggered based on increases in BOLD activation within the targeted ROI (the daIC in this example). **B)** Within each run of real-time fMRI-triggered experience sampling, participants viewed a visual fixation cross at the center of the screen. Presentation of the visual fixation cross was interrupted by trials of thought probes triggered by increases in BOLD activation in the run ROI, or by lack of brain-triggered trials after 100 seconds (“time-out” trials). Each experience sampling trial consisted of 8 thought probe items with 9-point Likert scales. Following the completion of an experience sampling trial, the visual fixation cross returned on the screen. C) For our first hypothesis, we predicted that subjective arousal ratings would be higher for trials triggered by daIC activation compared to other trial types. For our second hypothesis, we predicted that subjective external-attention ratings would be lower for trials triggered by PMC activation compared to other trial types.

## 2 Methods

### 2.1 Overview

The current study comprised two sessions taking place at the Temple University Brain Research and Imaging Center: (1) An initial 1-hour visit consisting of consent documentation, a study agenda overview, task training, and self-report questionnaires; and (2) The main 2.5-hour visit consisting of brief self-report questionnaires, task training recap, and 105 minutes of MRI procedures (see **Figure 1A**). During the MRI procedures, we collected localizer scans, fieldmap scans, and a T1-weighted anatomical scan prior to fMRI scanning. While undergoing fMRI scanning, participants completed an 8-minute run of the Gradual-onset Continuous Performance Task (GradCPT), which is our functional localizer task that was used to define ROIs within individuals (see section **“GradCPT”** for more information). Participants then underwent an 8-minute eyes-open resting state fMRI scan while we computed and visually inspected individual-level regions of interest (ROIs) from functional localizer data (see **Figure 1A)**. This resting state scan was not used in the final analysis. We then used the individual-level ROIs for our main experimental task, the rt-fMRI-ES paradigm (6 runs, 8-minutes each), in which experience sampling trials (see **Figure 1B**) were triggered based on BOLD activation level in a preselected ROI. The ROI for each rt-fMRI-ES run was selected with a counterbalanced order, yielding 3 runs targeting BOLD events in the daIC and 3 runs targeting BOLD events in the PMC.

### 2.2 Participants

Sixty adults were recruited from the larger communities of Drexel University and Temple University in Philadelphia, Pennsylvania, through study flyers and word of mouth. The ages of participants ranged from 18 years old to 35 years old (M=22.42, SD=4.29). Of the 60 participants, 24 identified as cisgender men, 34 identified as cisgender women, and one participant identified as a transgender woman. Racially and ethnically, participants self-identified as: White, Non-Hispanic or Latino (33.3%), White, Hispanic or Latino (8.3%), Black, Not Hispanic or Latino (16.6%), Black, Hispanic or Latino (1.6%), Asian, Not Hispanic or Latino (31.6%), and More than one race (8.3%). Participants were right-handed, fluent in English, free of any internal or external metallic objects contraindicative of a 3-Tesla magnetic field, had normal or corrected-to-normal vision, and had no history of a psychiatric, neurological, or chronic medical condition. Eligibility was assessed through a REDCap (Harris et al., 2009) pre-screening survey with questions about study-specific inclusion criteria and a subsequent telephone screener in which a study team member verbally reviewed an MRI Subject Safety Screening Form. Participants were paid for their participation at a rate of $35 USD per hour. All study protocols followed procedures that were approved by the Drexel University Institutional Review Board.

### 2.3 Procedures

#### 2.3.1 Day 1 Session

Following consent acquisition, participants began with study task training. During training for GradCPT, the task was colloquially described to the participant as the “Cities and Mountains Task.” Following instruction review, participants were presented with a 1-minute demo version of GradCPT to complete as practice. During task training for rt-fMRI-ES, the main objective of the task was framed as an attention task in which participants were to try to direct their attention to a fixation cross during the entirety of the run. Participants were informed that once every minute or so, the fixation cross would be replaced by a series of 8 questions that will ask about the content, quality, and dynamics of any thoughts that they were having directly preceding the appearance of the question set. To avoid performance bias, participants were not made aware of the time congruency of the thought probes relative to brain activity; timing of thought probe appearance was framed as being random. The definitions and scale anchors for each thought probe were thoroughly discussed. Following this, participants were instructed to complete a single-trial demo of the rt-fMRI-ES paradigm in which they were to rate responses to each question based on an example thought that had been read aloud by the researcher. This exercise was to gauge participant understanding of the task at hand and the meaning of each thought probe. Upon completing the demo, the researcher had participants verbally describe how they responded to each question and corrected any misunderstandings.

Participants privately completed a battery of mental health and behavioral questionnaires on REDCap in a testing room. This survey consisted of a section inquiring the participant’s legal full name and municipality of birth as it appears on their birth certificate to adhere to the National Institute of Mental Health National Data Archive Global Unique Identifier requirements (https://nda.nih.gov/). Questionnaires in this survey included the DSM-5-TR Cross-Cutting Symptom Measure – Adult (American Psychiatric Association, 2013), GAD-7 Anxiety (Spitzer et al., 2006), Patient Health Questionnaire 9 (Kroenke et al., 2001), Ruminative Response Scale (Nolen-Hoeksema & Morrow, 1991), STAI – Trait (Spielberger et al., 1983), World Health Organization Disability Assessment Schedule (Ustun et al., 2010), Perceived Stress Scale (Cohen et al., 1983), Adult ADHD Self-Report Scale (Kessler et al., 2005), Perseverative Thinking Questionnaire (Ehring et al., 2011), and the Mind Wandering Deliberate-Spontaneous Scale (Mrazek et al., 2015). Data from self-report questionnaires were not used in the final analysis.

#### 2.3.2 Day 2 Session

On the second visit of the study, participants began with a 45-minute preparation session where they completed an MRI Subject Safety Screening Form, a behavioral-state questionnaire, a review of the scanning agenda, and completion of a second review of task training for GradCPT and rt-fMRI-ES paradigms. The questionnaires consisted of the STAI – State (Spielberger et al., 1983) and the Stanford Sleepiness Scale (Hoddes et al., 1972) to assess the participant’s current state of anxiety and alertness. Metal checks were performed with a metal detecting wand and a ferromagnetic metal detecting pillar. The participant was set up on the MRI bed with the button box placed in their dominant hand to make responses during the tasks. An emergency squeeze ball was placed in the participant’s other hand in case they needed immediate attention during the session.

In the scanner control room, we first mapped the shared folder intended for incoming DICOMs (raw scanner images) on our project laptop to the Siemens console Y drive (see **“Real-time fMRI Analysis”** section). Once the folder was mounted, we used the *ideacmdtool* package (a tool that is installed on all Siemens consoles) to enable sharing of incoming images with whichever external drive was currently mapped. Within the shared folder, we created a subject folder for running our functional localizer package, rtfMRI-prep (see **“Functional Localizer Analysis”** section). We began the session with a localizer scan, Auto-Align scout scans, and an aligned localizer scan. Following the T1 image acquisition, we ran a B0 field map scan. Immediately following completion of the GradCPT fMRI task scan, our GradCPT MATLAB code output a text file that labeled the timing of each mountain event in the participant’s run. We transferred this text file to the subject folder created in the beginning of the session. In a Terminal window, we used *dcm2niix* (Li et al., 2016) to convert all existing DICOMs in the shared folder into NIFTI files. The NIFTI files of the participant’s T1-weighted structural image and GradCPT BOLD fMRI data were transferred into the subject folder. While the resting state fMRI scan was conducted, we ran our functional localizer package (see **“Functional Localizer Analysis”** section) within the subject folder containing the GradCPT event text file, the T1-weighted structural image, and the GradCPT BOLD fMRI image.

Once our functional localizer package finished running, we launched the Open Neurofeedback Training (OpenNFT, v1.0.0rc0) software to begin the first run of rt-fMRI-ES. OpenNFT is an open-source Python/MATLAB package that has the ability to detect incoming fMRI volumes, perform basic preprocessing, signal estimation and stimulus presentation for a variety of real-time fMRI protocols (Koush et al., 2017). In the OpenNFT graphic user interface, we loaded all necessary parameters that the software needs to perform processing and stimulus presentation during the run (see **“Real-time fMRI Analysis”** section). For efficiency, OpenNFT allows the user to input a configuration file that lists the desired settings of each necessary parameter. Our functional localizer package created these configuration files within the subject folder for each upcoming rt-fMRI-ES run. We then began the OpenNFT session, and started the rt-fMRI-ES scan. This process was repeated for each of the six rt-fMRI-ES task runs. To ensure task engagement, we actively checked for consistent and timely responses to each thought probe by observing the participant’s screen from a monitor in the control room during each run. We noted any signs of lack of engagement (i.e. lack of response to thought probes, which rarely occurred). In between each rt-fMRI-ES run, we verbally checked in with participants over an intercom to ensure that they were alert.

### 2.4 Neuroimaging Data Acquisition

Neuroimaging data were acquired using a Siemens 3-Tesla MAGNETOM Prisma MRI Scanner and a 64-channel head coil at the Temple University Brain Research and Imaging Center. Following the collection of localizer scans, a T1-weighted anatomical image was acquired with an MPRAGE sequence (repetition time: 2400 ms, echo time: 2.22 ms, inversion time: 1000 ms, flip angle: 8°, field of view: 256 x 256mm, 208 slices, 0.8mm isotropic voxels). We acquired a B0 field map for correction of echo-planar imaging (EPI) fMRI images (repetition time: 789 ms, echo time 1: 4.92 ms, echo time 2: 7.38 ms, flip angle: 45°, field of view: 192mm x 192mm, 2mm slice thickness, 2mm isotropic voxels). Functional magnetic resonance imaging (fMRI) runs were performed with a gradient EPI multiband T2* sequence (repetition time: 2000 ms, echo time: 25 ms, flip angle: 70°, field of view: 192mm x 192mm, 2mm slice thickness, 2mm isotropic voxels, multi-band acceleration factor: 3, phase encoding direction: anterior-to-posterior) with collection of 81 transverse slices aligned with the AC-PC plane. In total, 8 fMRI runs were collected per session for each participant (one GradCPT run, one resting state run, and six rt-fMRI-ES runs).

### 2.5 Experimental tasks

#### 2.5.1 GradCPT

The Gradual-Onset Continuous Performance Task (GradCPT) is a continuous performance task that engages sustained attention (Esterman et al., 2013). During a run of GradCPT, participants view a series of city- and mountain-scene grayscale images that quickly and continuously transition from one image to another every 800 ms (using linear pixel-by-pixel interpolation between consecutive images). Scene images are presented randomly, with city scenes frequently visualized (∼90% of total trials) and mountain scenes being displayed infrequently (∼10% of total trials), while the same scene cannot repeat on consecutive trials. Participants were instructed to press the index finger button on an MR-safe Current Designs 4-finger button box for every frequent trial (city scenes), and to withhold button presses for infrequent trials (mountain scenes). Performance can be assessed through analysis of reaction time, as well as accuracy.

Prior fMRI and intracranial electrophysiology studies have shown highly reliable and replicated salience network (including daIC) activation and DMN (including PMC) deactivation in response to infrequent stimuli (i.e., mountains) in the GradCPT (Esterman et al., 2013; Fortenbaugh et al., 2018; Kucyi et al., 2020; Kucyi & Parvizi, 2020). As such, GradCPT served as a functional localizer task to obtain the current study’s ROIs (daIC and PMC) within each individual participants’ native brain space. Participants completed a single 8-minute run of GradCPT, which was displayed using Psychophysics Toolbox in MATLAB (Brainard, 1997). Upon completion of GradCPT, the participant’s commission error rate (rate of erroneous button presses to infrequent mountain images) and omission error rate (rate of missed button presses during frequent city images) were calculated and displayed to the experimenter in the MATLAB Command Window. The GradCPT stimulus timing and fMRI data were used as input for an in-house functional localizer package, *rtfMRI-prep*, that generates individual-level regions of interest in daIC and PMC using a general linear model analysis (see **“Functional Localizer Analysis”** section). If the participant demonstrated inadequate performance effort during the task, as shown by a commission error rate of 0.75 or higher or an omission error rate of 0.33 or higher, an alternative script within *rtfMRI-prep* that generates standard space regions of interest rather than individual-level regions was to be used. While these GradCPT performance error rates are rather high (more than 5 times worse than the average performance in healthy adults; Esterman et al., 2013; Fortenbaugh et al., 2018), we intentionally set a liberal threshold as we were only trying to capture participants who were not performing at all, rather than individuals who had poor performance ability. While we developed an alternative, standard-space ROI option in place in the event of GradCPT performance being insufficient for use in functional localization, we did not have to use this method for any of the collected participants. All ROIs used during real-time analysis were functionally localized within each participant’s native brain space based on brain responses during GradCPT performance.

#### 2.5.2 Resting State

Following performance of the GradCPT, participants completed an 8-minute resting state fMRI scan. They viewed a central fixation cross and were instructed to keep their eyes open to prevent drowsiness. Participants did not make any task responses during this run. The fMRI data from this resting state run were not used for preregistered analyses. This scan served as an idle period to compute and visually inspect individual-level daIC and PMC ROIs (see section **“Functional Localizer Analysis”**), and to initialize the real-time fMRI analysis software that we used for our rt-fMRI-ES task runs (see section **“Real-time fMRI Analysis”**).

#### 2.5.3 Real-time fMRI-triggered experience sampling (rt-fMRI-ES)

We developed a novel experience-sampling paradigm, the real-time fMRI-triggered experience sampling (rt-fMRI-ES) task that we used to sample participants’ spontaneous mental experiences during six 8-minute fMRI runs. While this paradigm was used in conjunction with our real-time fMRI system to trigger thought probes based on brain activity, it was framed to participants as a visual fixation task with thought probes occurring randomly to prevent potential performance bias. In addition to triggering the appearance of the paradigm based on brain activity in either ROI, the paradigm appeared on the screen if no significant brain activation events had been detected by the real-time system after a predefined time window (termed “time-out” events / trials). Participants were instructed to focus their attention on a fixation crosshair and were informed that a set of 8 rating scales (a single trial) would appear at random times for them to report the qualities of thoughts that occurred immediately preceding the appearance of the trial (see **“Real-time fMRI Analysis”** for description of trial timing). Each rating scale was accompanied by a 9-point Likert scale with an arrow visualized on the center of the scale. Participants used an MR-safe Current Designs 4-finger button box to make responses. The middle finger button moved the arrow left, the ring finger button moved the arrow right, and the index finger button was used to submit participant responses to each scale. Only for the first rating scale of each set, there was an option to use the pinkie finger button to report that the participant was neither paying attention to the central fixation cross nor their internal mental thoughts. This option was included to distinguish thoughts that fall on some continuum of attention to task performance versus ongoing mental experience, from moments of external distraction or mind-blanking. Distraction/mind-blanking trials were excluded from analyses (see section **“Criteria for Excluding Data”**).

**Table 1** depicts the thought probes that comprised experience sampling trials. Rating scales were largely self-paced, with a 12-second time-out if response submission to each scale had not yet been made. Following the completion of each trial (a full set of the 8 rating scales), the visual fixation cross reappeared on the screen. Only thought probe items Q1 and Q3 (bolded in **Table 1**) were used for preregistered analyses (see section **“Statistical Analyses”**). The remaining thought probe items were collected for purposes that are beyond the scope of this registered report.

**Table 1.** Experience sampling thought probes. Eight thought probes comprise a single trial in the rt-fMRI-ES paradigm. The two probes used for our pre-registered hypothesis testing are in bold (Q1 External-Attention and Q3 Arousal).

| Thought Construct | Thought Probe | Low Anchor (“1”) | High Anchor (“9”) |
| --- | --- | --- | --- |
| **External-Attention** | **“I was paying more attention to the task than to my thoughts.”** | **Completely thoughts** | **Completely Task** |
| Affect | “I felt very positive.” | Extremely negative | Extremely positive |
| **Arousal** | **“I felt very activated / energized.”** | **Not at all** | **Completely** |
| Perceived repetitiveness | “This thought was similar to those I have had before.” | Not at all | Completely |
| Movement | “My thoughts were freely moving.” | Not at all | Completely |
| Engagement | “My thoughts were difficult to disengage from.” | Not at all | Completely |
| Rumination | “I was ruminating about the past.” | Not at all | Completely |
| Worry | “I was worrying about the future.” | Not at all | Completely |

## 3 Data Analyses

### 3.1 Functional Localizer Analysis (rtfMRI-prep)

We developed an in-house functional localizer package, *rtfMRI-prep* (https://github.com/DynamicBrainMind/rtfMRI_prep), which was used to identify the daIC and the PMC within the participants’ brains. The package was run promptly after collection of the T1-weighted anatomical scan and GradCPT fMRI scan, which were immediately accessible from our project laptop through real-time data sharing with the Siemens scanner console (see **“Day 2 Session”**). The script within *rtfMRI-prep* that we used during every neuroimaging session is *rtfmri_localizer_proc*, which identified the daIC and the PMC at the individual level based on participants’ BOLD activation during task-related events in the GradCPT. We developed an alternate script within the package, *rtfmri_localizer_proc_noGLM*, to use if a participant demonstrated inadequate performance effort during the GradCPT (i.e. a commission error rate of 0.75 or an omission error rate of 0.33; see section **“GradCPT”**). If run, this script generates regions of interest in standard space. As noted above, we did not use *rtfmri_localizer_proc_noGLM* for any of the collected participants. Using the participant’s anatomical scan, GradCPT fMRI scan, and a 3-column text file detailing GradCPT task event information, *rtfMRI-prep* performs basic fMRI preprocessing and computes functionally localized ROIs using a combination of FSL v6.0.7.7 (Jenkinson et al., 2012) functions in approximately 4.5 minutes on a high-powered Apple 14-inch Macbook Pro (Apple M3 Max Chip with 14 cores). The processes executed by the *rtfmri_localizer_proc* script within the *rtfMRI-prep* package are detailed below.

#### 3.1.1 Individual-level regions of interest (rtfmri_localizer_proc)

Using *rtfmri_localizer_proc*, motion correction was first performed on fMRI data using *MCFLIRT* (Jenkinson et al., 2002) with 6 transform degrees of freedom to align all volumes to the middle volume, followed by brain extraction of fMRI data using *BET2* (Jenkinson et al., 2005) with a fractional intensity threshold of 0.3. The anatomical image was skull stripped with *BET* (using robust brain center estimation), and we used *FLIRT* (Jenkinson & Smith, 2001) to perform linear registration (6 degrees of freedom, trilinear interpolation) between fMRI and anatomical images. We performed linear registration between T1 and MNI152 standard space using *FLIRT* (12 degrees of freedom, trilinear interpolation). Then, the obtained transformation matrices were concatenated to transform from the functional space to standard space. We immediately visually inspected the quality of image registrations using FSL’s *slices* tool. We then performed spatial smoothing of the brain-extracted functional image using *FSLMATHS* with a 6-mm full width at half maximum Gaussian kernel with mean filtering. As a final preprocessing step, we performed highpass filtering at 0.01 Hz using *FSLMATHS*.

Using the preprocessed fMRI data, we performed a general linear model (GLM) analysis using *FEAT* (Woolrich et al., 2001) with FMRIB’s improved linear model prewhitening. The GLM included one regressor for the onset of mountain (i.e., infrequent target) stimuli, convolved with a gamma hemodynamic response function (mean lag = 6 seconds; standard deviation = 3 seconds), plus the temporal derivative. We then searched for the top activated and deactivated voxels, respectively within the right daIC and bilateral PMC, based on voxelwise contrast of parameter estimate values. We constrained our search to voxels within standard (MNI152) space daIC and PMC masks obtained from publicly shared fMRI-derived functional parcellations that delineated these regions. For the daIC, we used cluster number 2 from a previously reported, data-driven 3-cluster insular cortex parcellation solution (Kelly et al., 2012). For the PMC, we obtained regions labelled as DMN within the Schaefer atlas (Schaefer et al., 2018) and included only DMN regions within the PMC cluster (which includes subregions of posterior cingulate cortex, precuneus, and retrosplenial cortex). Although we use the Schaefer atlas for the PMC due to clear delineation of this region within the DMN, we did not use the Schaefer atlas for defining the daIC because the relevant clusters in this atlas are contiguous with adjacent regions in the frontal operculum that are not the focus of our hypotheses.

We registered the daIC and PMC standard (MNI152) space masks to fMRI using the previously computed linear transforms. Subsequently, we retained the top 200 activated voxels (i.e., highest contrast of parameter estimate values) within the daIC mask and the top 200 deactivated voxels within the PMC mask to yield the final individual-level ROIs. Importantly, the middle fMRI volume from the GradCPT run, which was used for motion correction during functional localizer processing, was also used as a registration template for real-time fMRI analysis (as described in “Real-time fMRI Analysis”). As such, the individual-level ROIs were aligned with incoming fMRI data. After creation of individual-level ROIs, we immediately inspected functional anatomy of the daIC and PMC using FSL’s *fsleyes* tool.

### 3.2 Real-time fMRI Analysis (OpenNFT)

Real-time analysis of incoming fMRI volumes during runs of our rt-fMRI-ES task was conducted using OpenNFT v1.0.0rc0 software (Koush et al., 2017). OpenNFT has been successfully implemented in prior real-time fMRI studies to trigger neurofeedback and visual stimuli during behavioral performance (Pamplona et al., 2020). We used OpenNFT on a project-dedicated laptop (Apple 14-inch MacBook Pro with an Apple M3 Max chip with 14 cores) running Python 3.6.5. and MATLAB R2020b.

We facilitated data transfer between the MRI scanner and our project laptop via mapping a shared folder created on our laptop to the Siemens MRI console’s Y drive via TCP/IP connection. Once this connection was established, the scanner console deposited incoming DICOM files (single-frame data) into the mounted shared folder. Prior to starting each OpenNFT session (i.e., each run of rt-fMRI-ES), the file path of this shared folder was pre-specified in the OpenNFT session parameters as the folder that the software will monitor for incoming images (termed the “Watch Folder”). Other parameters that require pre-specification include the expected first file name, the repetition time, matrix size, and voxel size of brain volumes, and file paths of the folder containing the run’s ROI, the folder that OpenNFT will save data into following completion of the run, and the folders containing template functional and structural images of the participant’s brain.

Upon acquiring each individual brain volume, OpenNFT performed its default preprocessing steps using functions from SPM12 software (https://www.fil.ion.ucl.ac.uk/spm/), as described previously (Koush et al., 2017). These steps include spatial realignment to a template volume (from the GradCPT run; see above), estimation of six movement parameters (translation and rotation), reslicing, and spatial smoothing (5-mm full width at half maximum Gaussian kernel). Following OpenNFT default settings for further processing, we performed first-order auto-regressive correction to account for temporal autocorrelation that may be due to physiological noise (Lindquist, 2008). In order to account for global signal drift, we used several preprocessing tools. Specifically, we used an incremental GLM (iGLM) to regress out six head motion parameters and linear trends, and a high-pass filter was applied with a 1/128 sec cut-off. To further deal with high-frequency noise, a modified Kalman filter was applied (Koush et al., 2012).

For the current study, OpenNFT used an iGLM to calculate the BOLD percent-signal-change (%SC) of each incoming DICOM relative to the median BOLD baseline period, which was set to the first 10 brain volumes (20 seconds) of the run. After the baseline period, OpenNFT began to screen incoming data for candidate events. Though the run ROI (daIC or PMC) being screened for events was counterbalanced across all six runs of rt-fMRI-ES, OpenNFT screened activity in both ROIs for every run. In other words, while activity in both ROIs was monitored throughout the duration of each of the six runs, only one of the ROIs was the “target region” that rt-fMRI-ES stimuli (thought probe set) were triggered from for a given run. The second, nontarget ROI served as a control region. To trigger the appearance of rt-fMRI-ES stimuli based on BOLD activity in the target region of the given run, we used an algorithm that triggers stimulus presentation whenever the following criteria were met: (1) There was a consecutive BOLD %SC signal increase across three time points (volumes) in the target region; (2) At the current volume, the BOLD %SC of the target region was greater than the run’s baseline level (i.e., greater than zero value); and (3) Criteria #1 and #2 were not met in the non-target region; (4) At least 50 seconds (i.e., 25 TRs) have elapsed since the last thought probe set was triggered.

For the first criterion, it is important to note that not all three consecutive time points need to be high; in other words, we screened for evidence that activation in the target region was ramping up towards a peak. Our decision to trigger trials based on a successive increase across three time points in particular was motivated by simulations run on a pilot participant’s 8-minute resting data to compare the number of trials triggered when the algorithm searched for successive increases across a varying number of timepoints. The pilot participant’s resting state data was simulated using four versions of the triggering algorithm: a version that triggered a trial when there was an increase in brain activity across 2, 3, 4 and 5 TRs. Each of the four algorithms were simulated twice, one for each ROI (a run searching for activity in the daIC, and another run searching for activity in the PMC). While the versions that triggered trials based on increases across 4 TRs and 5 TRs were overly conservative (4 TRs triggered 4 PMC-triggered trials and 5 daIC trials; 5 TRs triggered 3 PMC trials and no daIC trials), the version that triggered trials based on increases across 2 TRs proved to be excessively liberal by triggering the maximum number of trials possible in both brain regions (7 trials in both runs). Thus, the algorithm that triggered trials based on a successive increase in brain activity across 3 TRs was the most reasonable option (7 PMC-triggered trials, 6 daIC-triggered trials). Moreover, upon inspecting the events that were triggered by the different algorithms, we found that the 3-TR algorithm, relative to the other algorithms, more reliably detected events that preceded or were at BOLD peaks.

Importantly, even when accounting for both the 6-second delay that is consequential of the first criterion and the ∼4-6 second hemodynamic delay characteristic of BOLD activity measurements, the timing of thought probe onset in the current study was more efficient in capturing significant activity associated with mind wandering compared to traditional, randomly timed experience sampling methods. Our past research shows that significant BOLD activity in DMN subregions is often present and sustained as early as ∼20 seconds prior to thought probe onset when using random-interval experience sampling (Kucyi et al., 2024). Therefore, though a thought probe in the current study was likely displayed about 12 seconds following the initial arising of a neural event of interest, this was likely far timelier in capturing this activity than typical random-onset experience sampling methods.

We selected our algorithm criteria based on simulations of previously collected fMRI data using OpenNFT’s offline session feature. By monitoring activity in both ROIs, we can ensure that thought probe sets triggered by brain activity in the target region would not otherwise be triggered by the non-target region. This is vital to guarantee that the neural events triggered by the two ROIs are distinct. It is important to note that our decision to concurrently screen activity in both ROIs was made after receiving peer-review feedback on our Stage 1 preregistration report, which was after the first 15 participants were collected. During the rt-fMRI-ES runs of the first 15 participants, the OpenNFT software was configured to only monitor activity in the target region of each run. After changing our algorithm to screen both regions, we retroactively simulated our real-time fMRI protocol for the first 15 participants’ runs to see if any of their completed trials would have also been triggered by the non-target region. Across all participants’ total completed trials, an average of 3 trials per participant would have also been triggered by non-target regions. We excluded these trials from the final analysis. Furthermore, the exclusion criteria ensured that thought probe sets did not appear with excessive frequency and that triggers were initiated when the slope of BOLD %SC in the target region was increasing, thereby signifying the onset of an ‘activation’ event. If no significant increase was detected after a considerable period of time since onset of the last thought probe set (100 seconds), then the experience sampling paradigm was automatically triggered (termed “time-out” events / trials). We used Psychophysics Toolbox in MATLAB R2020b to present our visual stimuli (Brainard, 1997) (see section **“Real-time fMRI-triggered experience sampling”**).

### 3.3 Statistical Analyses

The hypotheses that we tested in this study involved analyzing experience-sampling ratings as a function of trial type (i.e., how an experience-sampling probe was triggered). Because trial type depended on real-time fMRI analysis, the number of observations for each trial type was unbalanced across participants. We consequently chose to test both hypotheses using linear mixed-effects models, which are particularly well-suited for estimating effects in designs where observations from participants are unbalanced across an experimental design (Pinheiro & Bates, 2000). All mixed effects models were fit using the *lme4* package (Bates et al., 2015) in the *R* statistical software (R Core Team, 2022).

To test our first set of hypotheses (*H1*), **that subjective ratings of arousal time-locked to daIC-activation trials will be higher than ratings time-locked to non-daIC-activation trials (including both PMC-activation trials and time-out trials),** we modeled subjective ratings of arousal (response to Q3 prompt, “I felt very activated / energized.”) as a function of trial type. Trial type was included in the model as a dummy-coded categorical fixed effect with two levels: (1) daIC-activation triggered probe (coded as 1); and, (2) non-daIC-activation triggered probe (coded as 0). Note that level (2) encompasses both PMC-activation triggered probes and time-out triggered probes, which were grouped together because time-out triggered probes were sparse, and both categories are defined by a lack of daIC activation. This dummy coded variable yielded one slope parameter capturing the change in subjective ratings of arousal between daIC-activation triggered probes and non-daIC-activation triggered probes. We included trial-wise head motion, estimated based on mean relative displacement estimated with FLIRT for the 5 TRs prior to thought probe onset, as a covariate to control for (see section **“Criteria for Excluding Data”**). To account for between-participant variability in the fixed effects, we included the slope contrasting activation conditions, the model intercept, and all covariances between the fixed effects, as random effects grouped by participant. In doing so, we specify the maximal random effects structure allowed by the data, and we iteratively simplified the model by dropping random effect parameters until (1) the model reaches convergence, and (2) the dropped random effect parameters prove to be significant based on likelihood ratio tests (LRTs; following prior recommendations (Barr et al., 2013)). We predicted that the slope contrasting daIC-activation against PMC-activation and time-out trials would be significantly positive, suggesting that daIC-activation triggered trials are associated with higher levels of subjective arousal ratings relative to the other two trial types. We estimated *p*-values for significance tests from well-established simulation methods using *lmerTest* in *R* (i.e., the Satterthwaite degrees of freedom approximation; (Kuznetsova et al., 2017)), since assuming the null distribution of fixed effects parameters follows standard *t* or *F* distributions is often violated in practice (especially for unbalanced designs), and LRTs are known to produce anti-conservative *p*-value estimates (Pinheiro & Bates, 2000).

To test our second hypothesis (*H2*), **that subjective ratings of external-attention time-locked to PMC-activation trials will be lower than ratings time-locked to non-PMC-activation trials (including both daIC-activation trials and time-out trials)**, we conducted a similar linear mixed effects model as that reported above. However, instead of self-reported arousal, self-report external-attention (response to Q1 prompt, “I was paying more attention to the task than my thoughts”) was modeled as a function of trial type. Trial type was again a dummy coded fixed effects categorical factor with two levels: PMC-activation triggered probe (coded as 1), and non-PMC-activation triggered probe (coded as 0). Similar to the previous hypothesis, non-PMC-activation triggered probe encompassed both daIC-activation triggered probes as well as time-out triggered probes. We specified the model using the same methods described above: by specifying the maximal random effects structure allowed by the data and iteratively simplifying the model until convergence is reached and until random effects factors prove to be significant based on LRTs (Barr et al., 2013). We predicted that the slope contrasting PMC-activation against daIC-activation and time-out trials (*H2*) will be significantly negative, suggesting that PMC-activation triggered probe trials are associated with lower levels of self-reported external-attention relative to the other two trial types. All *p*-values were estimated using the same methods described above for testing our first hypothesis.

### 3.4 Power Analysis

The proposed sample size of 60 was determined by conducting a power analysis on data from a previous fMRI study from our group that used random-interval experience sampling to assess mind-wandering (defined as task-unrelated thought, similar to Q1 in the current study) (Kucyi et al., 2016). These data consisted of 36 thought probes per participant, with DMN activity estimated from preprobe periods, for each of 28 participants. We assumed that the effect sizes we will observe in the present study (i.e., between activity in the ROIs and reports on fMRI-triggered experience sampling) would be at least as strong as that observed in this previous study. As noted, however, our rt-fMRI-ES paradigm is designed to enhance statistical power relative to random-interval experience sampling. Moreover, we used functional localizers to target individual-specific ROIs during rt-fMRI-ES, a procedure that intends to enhance statistical power relative to prior work. As such, our power calculation may be viewed as conservative.

Data from the present study was analyzed with a linear mixed model, and so, for consistency, we fit a linear mixed model to data from the previous study to estimate how many participants would be needed to achieve sufficient power. We modeled mind-wandering reports (0 to 100 scale with 100 being the most mind wandering) as a function of DMN percent signal change and included a random intercept and slope grouped by participant using the *lme4* package (Bates et al., 2015) in R (R Core Team, 2022). The coefficient estimate was 23.425, suggesting that each unit increase in DMN percent signal change was associated with a 23.425 unit increase in self-reported mind wandering (*b =* 23.425, 95% CIs = [9.046, 39.964], *t*(19,561) = 3.058, *p* = .006). We used the fitted model to estimate power across a range of possible sample sizes using Monte Carlo simulations (1000 simulations each for sample sizes ranging from 3 to 45 in steps of roughly 5) using the *powerCurve* function from the *simr* package (Green & MacLeod, 2016) with an alpha level set to 0.02. These simulations revealed that a sample size of 40 resulted in power levels of 0.857, sufficiently beyond the conventional 0.80 power threshold (*M* = .857, *95 CI* = [.834, .878]). To be conservative, we recruited 20 extra participants to account for participants who may fail to meet our criteria for inclusion in the study. This yielded a total recruitment number of 60 participants.

## 4 Criteria for Excluding Data

### 4.1 Preregistered Criteria for Trial-level Exclusion

Following a review of the final collected dataset, we excluded data based on preregistered criteria. We excluded data due to excessive head motion, which could invalidate rt-fMRI analysis estimates and lead to erroneously triggered experience-sampling prompts. Head motion was assessed through both visual inspection of the fMRI images using the FSL tool *FSLeyes* (McCarthy, 2021), and through motion estimates using FSL’s *MCFLIRT* (Jenkinson et al., 2002). Following prior recommendations (Power et al., 2014), we used frame-wise displacement (FD) as the main metric to be used for excluding trials and/or participants. Furthermore, we excluded any trials that were presented soon after (<10 seconds) the completion of the previous thought probe set. While our algorithm for triggering trials based on high-activation brain states was carefully written to mitigate quick consecutive trial onset (see section **“Real-time fMRI Analysis”**), a trial may have occasionally followed another quickly after completion if the participant took a long time to answer each thought probe throughout the trial. If a trial quickly followed another, the participant likely did not have an ample amount of time to re-engage in attentional fluctuations toward and away from visual fixation. Furthermore, if a participant pressed the pinkie button for the first thought probe of a trial indicating they were neither focused on the fixation cross nor their internal mental thoughts (i.e. distracted by external stimuli, or their mind was blank), we excluded this trial. To account for lack of alertness during a trial, we excluded any trials for which a participant recorded ratings for less than half of the eight thought probes. As described in section **“Real-time fMRI Analysis (OpenNFT)”**, our real-time triggering algorithm was configured to track BOLD activity in both the target and non-target region in each run of rt-fMRI-ES following reviewer feedback on our Stage 1 registered report after 15 participants had already completed the full study protocol. Prior to this implementation, trials were triggered based on increasing activation in the target region without accounting for BOLD activation in the non-target region. Thus, we retroactively simulated our real-time fMRI protocol for the first 15 participants, and we excluded completed trials that would have also been triggered by the non-target region. In summary, we excluded individual trials when the following criteria were met:

- The presence of any volume with an FD value greater than 0.25 mm across the five volumes prior to onset of thought probes. Even though we used data from three volumes to screen for events (see section **“Real-time fMRI Analysis”**), it is known that large head movements can impact subsequent frames (Power et al., 2014), and so we performed exclusion based on five pre-probe volumes as a conservative measure.
- The trial appeared less than 10 seconds after the completion of a previous trial.
- For item 1 of the trial (Q1; see section **“Real-time fMRI-triggered experience sampling (rt-fMRI-ES)”**), the participant indicated they were neither focused on the fixation cross nor their internal mental thoughts.
- The participant recorded ratings for less than half of the thought probes in that trial.
- For the first 15 participants, the trial would have also been triggered by increasing BOLD activation in the non-target region (as revealed by retroactive computational simulations of the participant’s data).

### 4.2 Preregistered Criteria for Subject-level exclusion

Additionally, we excluded all data from a participant when the following criteria were met:

- More than half of the participant’s trials were excluded due to excessive head motion (based on trial-wise criteria specified above).
- More than 75% of the participant’s trials were “time-out” events not triggered by BOLD activation (see section **“Real-time fMRI-triggered experience sampling”**).
- During more than 50% of the participant’s trials, they were neither focused on the fixation cross nor their internal mental thoughts (see section **“Real-time fMRI-triggered experience sampling (rt-fMRI-ES)”**).
- During data acquisition, the participant was unable to complete more than 4 runs of rt-fMRI-ES.

### 4.3 Preregistered Criteria for Statistical Model Exclusion

To retain power for both of our proposed statistical tests (see section **“Statistical Analyses”**), we imposed a minimum for both brain-triggered trial types (daIC-activation trials and PMC-activation trials) at 8 total observations for all participants. Thus, the following exclusion criteria applied to all participant data for either statistical test:

- A participant was excluded from the model testing the daIC hypothesis (H1) and not from the model testing the PMC hypothesis (H2) if the participant completed fewer than 8 daIC-activation triggered trials.
- A participant was excluded from the model testing the PMC hypothesis (H2) and not from the model testing the daIC hypothesis (H1) if the participant completed fewer than 8 PMC-activation triggered trials.

We did not recruit new participants in place of those excluded post-data collection. Based on our power analysis, we concluded that a minimum sample size of 40 would be sufficient in retaining power for our statistical analyses (see section **“Power Analysis”**).

## 5 Results

### 5.1 Description of Final Dataset

Of the 60 participants who completed the full protocol, two were excluded prior to trial exclusion analyses due to technical errors during their neuroimaging sessions. Trial-level data from the remaining 58 participants were evaluated according to the preregistered exclusion criteria listed in *Criteria for Excluding Data*. **Figure 2A** illustrates the proportion of trials that were excluded for each participant broken down by exclusion reason. Seven participants did not require any of their trials to be excluded. Among the first 15 participants, an average of three trials would have also been triggered by increasing BOLD activation in the non-target region (blue segments labeled “Opposite ROI-triggered” in **Figure 2A**). While all data from the first two participants in this initial cohort were excluded prior to final analyses, “Opposite-ROI triggered” exclusions were not solely responsible for these participants falling below the minimum trial inclusion criteria for analyses. For seven participants (including the first two participants of the initial recruitment cohort), over half of their total trials were discarded (participants with less than 20 remaining trials in **Figure 2C**); we excluded all data from these seven participants prior to final analyses due to an insufficient number of remaining trials.

**Figure 2.**
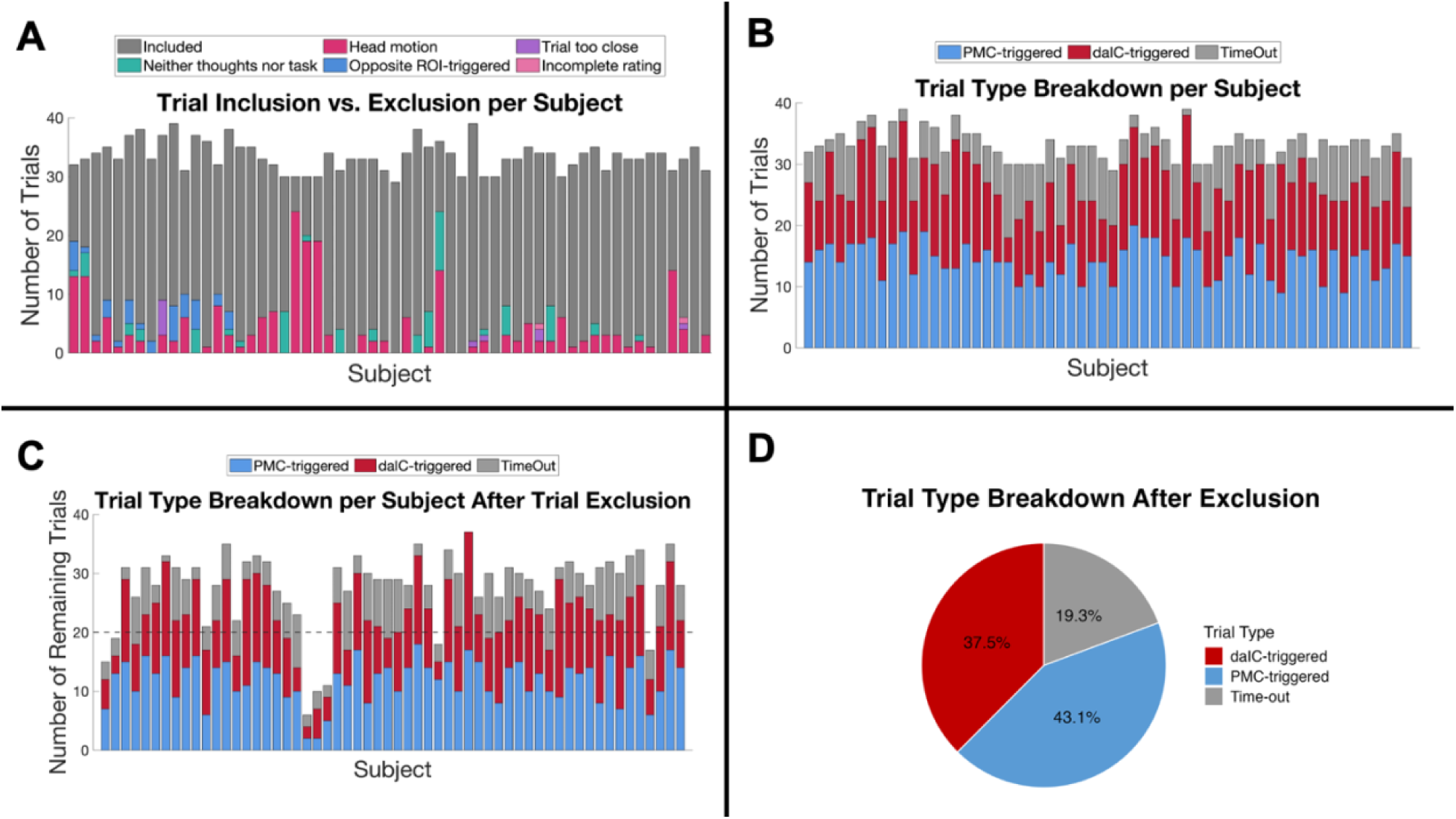
Summary of Trial Exclusions. **A)** Trials were excluded from each participant’s data for the following reasons: a trial had excess head motion (“Head Motion” – hot pink), a trial followed too close after another (“Trial too close” - purple), the participant reported that they were neither focused on their thoughts nor the task for Q1 of the trial (“Neither thoughts nor task” - green), the trial would have been triggered by the non-target region of interest (“Opposite ROI-triggered – blue), and the participant answered less than half of the rating scales in the trial (“Incomplete rating” – light pink). Retained trials are represented in gray. **B)** Subject-level breakdown of the proportion of trial types that were collected during the real-time session before trial exclusion procedures. daIC-triggered trials are in red, PMC-triggered trials are in blue, and time-out trials are in gray. **C)** Subject-level breakdown of the proportion of trial types that were considered for final analysis after trial exclusion procedures. Color labels are the same as 2B. **D)** Group-level breakdown of the proportion of trial types considered for final analysis following trial exclusion procedures. Color labels are the same as **2B** and **2C**.

Prior to the trial exclusion procedures described above, an average of 12.67 trials that were triggered by increased daIC activation (SD=3.47) was observed per participant (**Figure 2B**). The mean number of viable daIC-triggered trials across participants reduced to 10.4 (SD=4.13) following trial exclusion (**Figure 2C**). On average, participants completed 14.36 trials (SD=2.91) that were triggered by increased PMC activation. The mean number of viable PMC-triggered trials across participants reduced to 11.96 (SD=3.96) following trial exclusion (**Figure 2C**). In total, the remaining trials across all participants were divided as follows: daIC-triggered (37.5%), PMC-triggered (43.1%), and time-out events (19.3%; **Figure 2D**). Taken together, with 51 participants eligible for final analyses, we retained enough data to meet our pre-registered power analysis requirement (see **“Power Analysis”**).

### 5.2 Exploratory Validation of Real-Time BOLD Signal Triggers

Interpretations of the present study’s results are grounded in the assumption that our real-time BOLD signal estimates (i.e., obtained online) are similar to those procured through typical fMRI processing procedures after the completion of neuroimaging session (i.e., obtained offline, typically with more intensive computing for artifact removal). Thus, we conducted exploratory validation analyses to compare the alignment between BOLD signal values obtained online versus offline at time points surrounding trigger events. We focused our visual validation on BOLD activity in the 32-second time windows centered on trigger times for daIC-activation and PMC-activation trials that surpassed inclusion criteria. Importantly, as we were constrained to our pre-registered analysis, our validation was solely visual in nature and did not involve statistical or quantitative comparisons of the time series data. The goals of this exploratory validation were (1) to ensure that, for either ROI, the general trends and shapes of the time series obtained in real-time were similar to those of the offline preprocessed time series; and (2) to assess whether, for both real-time and offline time series, target and non-target BOLD activity near either brain-triggered trial type was congruent with region activity expectations for that trial type. In other words, for trials that were triggered by daIC activation, we would expect to see both the real-time and offline daIC time series to increase leading up to trial onset, and both the real-time and offline PMC time series to not increase prior to trial onset.

Real-time fMRI preprocessing and region time series acquisition steps are described above under section **“Real-time fMRI Analysis (OpenNFT)”**. Offline preprocessing steps involved Independent Component Analysis Automatic Removal of Motion Artifacts (ICA-AROMA), a rigorous and computationally intensive method for motion artifact removal (Pruim et al., 2015). We applied ICA-AROMA along with a preprocessing pipeline used in our past work, as detailed elsewhere (Kucyi et al., 2016, 2024). To obtain offline region time series, we extracted mean time series values of either ROI from each run’s preprocessed fMRI data using subject-specific region masks created in-session (see **“Functional Localizer Analysis (rtfMRI-prep)”**). The validation involved comparison of four time series types across all runs completed by 51 subjects included in final analyses: (1) real-time daIC; (2) offline daIC; (3) real-time PMC; and (4) offline PMC. For each time series type, data were *z*-scored within subjects to account for differences in value ranges between real-time and offline data. We extracted 32-second epochs around both daIC-triggered trials and PMC-triggered trials from each time series. For either event validation (daIC-triggered and PMC-triggered), epochs for each of the four time series types were averaged within subjects. We calculated the group-level grand average of within-subject averaged epochs for each time series, yielding four time series per event validation.

Figure 3A shows average estimates of real-time daIC activity (target region), offline daIC activity, real-time PMC activity (non-target region), and offline PMC activity near the onset of daIC-triggered trials. Both real-time and offline estimates of daIC activity steadily increase in the 8-second pre-probe period. Estimates of real-time and offline PMC activity sustain a flat shape throughout the time window, although a delayed decrease is notable after trial onset (in line with an expected deactivation that is characteristic of the DMN).

**Figure 3.**
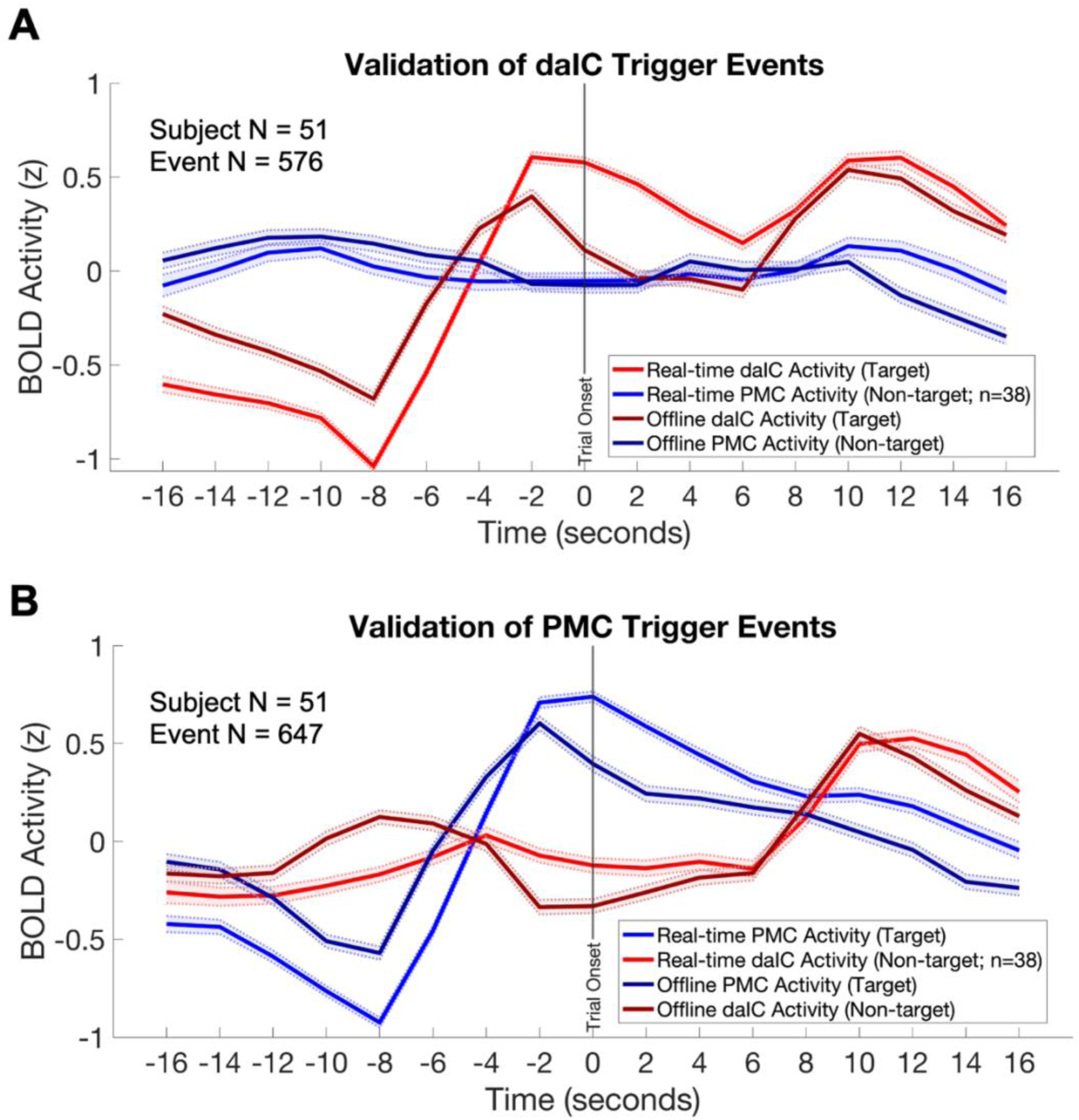
Exploratory Validation of Real-Time BOLD Activity Triggers. **A)** daIC Trigger Events: Each line comprises data averaged across subjects after computing within-subject averages for the respective data type (as described in the figure legend ). Shaded error bars depict the standard error of the mean (SEM) of across-subject activity at each time point. Across all trials that were triggered by increasing daIC activity in real-time, on average, real-time daIC activity estimates increased over the eight seconds leading up to the trial onset. This pattern was also found in daIC activity estimated from more traditional offline preprocessing measures. The time series of real-time PMC activity during daIC-triggered trials only contains data from 38 subjects, as the first several subjects of the study underwent rt-fMRI-ES prior to the implementation of tracking real-time activity in the non-target region. **B)** PMC Trigger Events: The same data averaging procedures and SEM calculations as **3A** were conducted. Across all trials that were triggered by increasing PMC activity in real-time, on average, real-time PMC activity estimates increased over the eight seconds leading up to the trial onset. This pattern was also found in PMC activity estimated from more traditional offline preprocessing measures. Similar to **3A**, only 38 subjects had real-time data that estimated activation in the non-target daIC region.

Figure 3B shows average estimates of real-time PMC activity (target region), offline PMC activity, real-time daIC activity, and offline daIC activity near the onset of PMC-triggered trials. Both real-time and offline estimates of PMC activity increase in the 8-second period prior to trial onset. Notably, estimates of real-time and offline daIC activity remain relatively flat between the 16-second pre-probe to 6-second post-probe period, then increase about 8-seconds after trial onset. A similar trend of real-time and offline daIC activity increasing 8-seconds past the start of the trial is observed in Figure 3A. This likely reflects an increase in daIC activity during the action of performing thought probe ratings, which is in line with the daIC’s role in effort-based task performance (Aben et al., 2020; Nelson et al., 2010; Touroutoglou et al., 2012).

Overall, we confirmed that our real-time estimates of BOLD activation in each ROI were largely similar to typical offline measures of ‘cleaned up’ BOLD signal. Furthermore, for both real-time and offline estimates of either ROI, we visually noted expected BOLD activity trends around both daIC trigger events and PMC trigger events. Through validating our real-time BOLD signal estimates, we can verify that our preregistered results are based on BOLD estimates that are similar to those derived from traditional methods of brain signal acquisition.

### 5.3 Hypothesis 1 Testing: Arousal ratings during daIC-triggered trials

Out of the remaining 51 participants who survived subject-level exclusion, 42 participants were eligible for inclusion in the linear mixed-effects model testing our first hypothesis that ratings of arousal would be higher for trials time-locked to increased daIC activation compared to other trial types. Ten participants were excluded from the model due to an insufficient number of daIC-triggered trials following trial-level exclusion (less than eight daIC-triggered trials; see **“Statistical Model Exclusion”** and Figure 2C). A total of 1243 trials across all eligible participants were included in the model.

The model showed that there was no significant effect of triggering trials based on increased daIC activity on arousal ratings (Figure 4A**, top**) (*b* = –0.02, 95% CI [-0.25, 0.20], df = 39.7, t = –0.21, p = .84). While arousal ratings for daIC-triggered trials were slightly lower on average compared to arousal ratings during non-daIC-triggered trials (such as PMC-triggered and time-out trials), trending in opposition to our hypothesis, this effect did not reach statistical significance.

**Figure 4.**
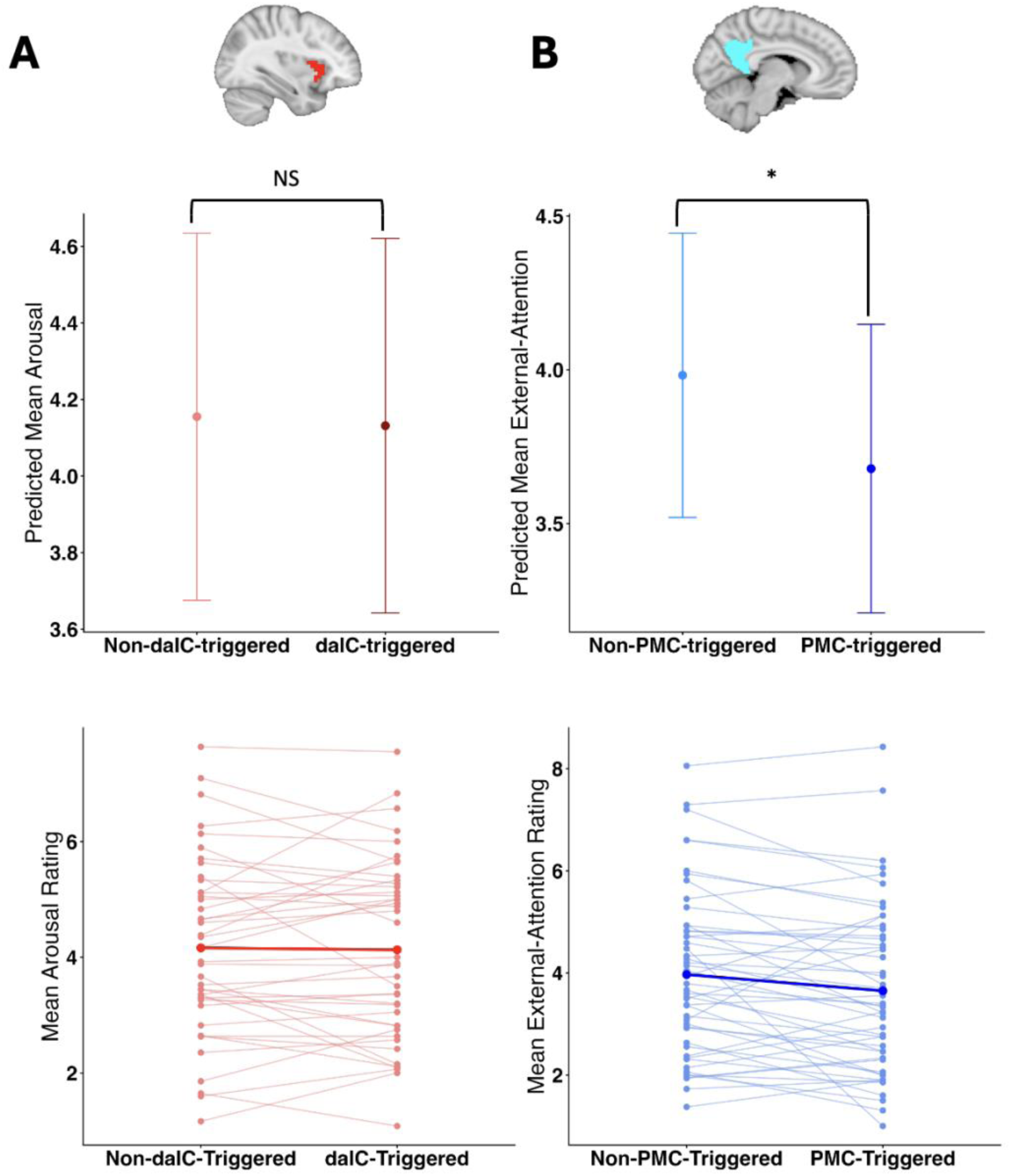
Results of Main Statistical Analyses. **a)** daIC and Arousal (H1): There was no significant difference between model-estimated mean arousal during trials triggered by the daIC versus non-daIC-triggered trials. Error bars reflect approximate 95% confidence intervals around the predicted mean. In the paired plot, each thin pink line connects a single subject’s mean arousal during non-daIC-triggered trials to their mean arousal during daIC-triggered trials. The bold red line connects the group average arousal during non-daIC-triggered trials to average arousal during daIC-triggered trials. **b)** PMC and External-Attention (H2): There was a significant difference between model-estimated mean external-attention during trials triggered by the PMC and non-PMC-triggered trials. Error bars reflect approximate 95% confidence intervals around the predicted mean. In the paired plot, each thin light blue line connects a single subject’s mean external-attention during non-PMC-triggered trials to their mean external-attention during PMC-triggered trials. The bold dark blue line connects the group average external-attention during non-PMC-triggered trials to average external-attention during PMC-triggered trials.

As an exploratory test of whether the order of target region selection influenced the effect of triggering from the daIC on arousal ratings, we included counterbalance order as a categorical dummy-coded interaction term in the model with two levels: (1) the daIC was the target region for every odd-numbered run (coded as 1), and (2) the PMC was the target region for every odd-numbered run (coded as 0). This exploratory model demonstrated that there was no significant interaction between daIC-triggering and the counterbalance order of target region selection when estimating arousal ratings (*b* = -0.07, 95% CI [-0.52, 0.38], df = 38.8, t = -0.29, p = .77).

When visualizing how average arousal differed between trial types by subject, 20 out of the 42 subjects (47.6%) included in the model had higher mean arousal scores during daIC-triggered trials compared to non-daIC-triggered trials, trending in the hypothesized direction (Figure 4A**, bottom**). Twenty-one subjects had lower mean arousal scores during daIC-triggered trials compared to non-daIC-triggered trials. One subject had equivalent mean arousal scores for both conditions.

### 5.4 Hypothesis 2 Testing: External-attention ratings during PMC-triggered trials

Forty-nine of the remaining 51 participants were included in the model testing our second hypothesis. Three participants had an insufficient number of PMC-triggered trials for the model following trial-level exclusion (less than eight PMC-triggered trials; see Figure 3C). A total of 1429 trials across all eligible participants were included in the model.

The model revealed a significant negative effect of PMC triggering on external-attention ratings (Figure 4B**, top**) (*b* = –0.30, 95% CI [-0.55, -0.06], df = 45, t = –2.50, p = .016). In support of our hypothesis, PMC-triggered trials were significantly associated with lower external-attention ratings compared to trials that were not triggered by the PMC. On average, external-attention ratings were lower by 0.30 of a unit for trials time-locked to PMC activation compared to other trial types.

Similar to the exploratory model described above in **“Hypothesis 1 Testing: Arousal ratings during daIC-triggered trials”**, we performed an exploratory analysis to examine whether counterbalance order of the selected target region influenced the relationship between triggering trials from increasing PMC activation and ratings of external-attention. The model showed that there was no significant interaction between PMC-triggering and the counterbalance order of target region selection when estimating external-attention ratings (*b* = -0.18, 95% CI [-0.66, 0.30], df = 43.7, t = -0.74, p = .47).

When visualizing how average external-attention differed between trial types by subject, 33 out of the 49 subjects (67.3%) included in the model had lower mean external-attention scores during PMC-triggered trials compared to non-PMC-triggered trials, trending in the hypothesized direction (Figure 4B**, bottom**). Sixteen subjects had higher mean external-attention scores during PMC-triggered trial compared to non-PMC-triggered trials.

## 6 Discussion

### 6.1 Overview

To our knowledge, the present study is the first to develop and validate real-time fMRI-triggered experience sampling (rt-fMRI-ES). We tested two hypotheses: (1) increased activation of the daIC would be time-locked to thoughts that are higher in arousal (a theoretically novel examination); and (2) increased activation of the PMC would coincide with mental experiences that are lower in external-attention (a replication of an established brain-behavior relationship). We did not find support for our first hypothesis, as there was no significant difference in arousal ratings between trials that were triggered by daIC activation and other trial types. However, external-attention ratings were significantly lower during trials that were triggered by PMC activation than other trial types, providing evidence for our second hypothesis. Although our speculative daIC hypothesis was not supported, our finding of the classic relationship between the PMC and stimulus-independent thought provides proof-of-concept for rt-fMRI-ES as a method to study the neural basis of ongoing thought.

### 6.2 The daIC and Subjective Arousal

Often considered a part of the brain’s salience or cingulo-opercular network (Seeley et al., 2007; Uddin et al., 2023), the daIC has been implicated in myriad cognitive processes and has been shown to activate across multiple task domains. For example, the daIC is involved in cognitive control and the detection of novel stimuli (Uddin et al., 2017) as well as ongoing self-monitoring of task-based errors (Ullsperger et al., 2010). Additionally, spontaneous activity in the daIC has been shown to correlate with neurophysiological markers of arousal during wakeful rest (Kucyi & Parvizi, 2020). Thus, while activity in the daIC has been associated with a variety of seemingly disparate experimental operations, one possibility is that its role in may relate to increased general arousal.

Arousal has been operationalized using a range of outcome measures, lending to its rather ambiguous conceptualization in the literature (Reid et al., 2025; Smith et al., 2025). A recent meta-analysis of neuroimaging studies on arousal identified several “forms” of arousal, including cognitive arousal, affective arousal, and physiological arousal (Sabat et al., 2025). While each form of arousal possessed distinct neural representations, activity in the daIC was found to be correlated with a variety of arousal types. In other words, the daIC has been considered as a host for multiple dimensions of arousal. However, causal evidence for a role of the daIC in arousal has not been conclusive. For example, while intracranial electrophysiological recordings from the daIC have demonstrated increased high frequency activity during a sustained attention task compared to other areas of the insula, direct stimulation of this area failed to consistently evoke changes in subjective experience (Duong et al., 2023).

Importantly, the neural correlates of arousal have typically been studied within task-based contexts rather than a wakeful resting state as applied in our rt-fMRI-ES paradigm. Additionally, our measurement of arousal as subjective self-reported “activation” differs from commonly used physiological arousal measures. Past work from our group examined the physiological associations of self-reported arousal during a random-onset experience sampling task performed during electroencephalogram (EEG) recording (Braun et al., 2025). Notably, fluctuations in self-reported arousal showed minimal correlation to multiple physiological measures commonly linked to arousal, including EEG alpha power and heart rate variability (Braun et al., 2025). This finding is in line with past research reporting an inconsistent relationship between subjective self-reported arousal and physiological arousal (Agren, 2023; Leonidou & Panayiotou, 2022; Wenzler et al., 2017). Taken together, the present study’s findings (i.e., a lack of relationship between daIC activation and subjective arousal at rest) build on accumulating evidence that subjective arousal likely differs fundamentally from traditional task-evoked “objective” arousal.

An additional consideration for our finding is the possibility that different participants performed ratings with differing interpretations of the arousal probe (“I felt very activated / energized”). Each participant received thorough training on the conceptual definitions of the thought probes prior to rt-fMRI-ES and were instructed to consider their level of affective energy in relation to their thoughts for the arousal probe. Despite this, participants could have been evaluating their mental experiences with varying understandings of arousal as a construct, such as conflating arousal with wakefulness. Thus, it is possible that our arousal probe, intended to capture affective arousal, inadvertently also sampled multiple other dimensions of arousal, a common practical challenge with self-reported experience sampling measures (Shareef-Trudeau et al., 2025).

### 6.3 The PMC and External Attention

Activation in the posteromedial cortex (PMC; comprising the precuneus and the posterior cingulate cortex) has consistently been shown to coincide with both stimulus-independent rest and tasks that involve the self-generation of mental content (Andrews-Hanna, 2012; Christoff et al., 2009; Spreng et al., 2009). Thought to be a core region of the DMN (Buckner & DiNicola, 2019), the PMC exhibits functional connectivity to other DMN nodes at rest and displays decreased activity during engagement with externally-oriented goal-directed actions (Cavanna & Trimble, 2006; Utevsky et al., 2014). Extant research using random-onset experience sampling during fMRI shows that increased DMN activation correlates with self-reported ratings of mind-wandering and off-task thought (Christoff et al., 2009; Kucyi et al., 2013, 2016; Mittner et al., 2014; Stawarczyk et al., 2011). Across various contexts, the role of the PMC has be characterized as the central locus of self-referential internal mentation (Cavanna & Trimble, 2006).

In line with past research, we observed significant decreases in self-reported external-attention during experience sampling trials that were triggered by spontaneous PMC activation compared to other trial types. Our findings offer a novel perspective to this established relationship by demonstrating a link *from* increases in ongoing PMC activity *to* decreased external-attention, an inverse of how brain-behavior relationships are classically identified. The ability of the rt-fMRI-ES method to capture moments of lower external-attention as they unfold positions rt-fMRI-ES as a promising tool for efficiently studying the theorized neural substrates of ongoing thoughts.

Taken together, our results are not surprising; the PMC’s role in internally-oriented attention is well established, whereas the relationship between the daIC and subjective arousal may be more complex and multidimensional. The connection between the daIC and reportable experience may be less direct, or may be more obvious in stimulus-evoked contexts that differ from the current study’s parameters. While our self-report measure of attention orientation essentially evaluates the degree to which an internal mode of mentation is present, our subjective arousal probe gauges a specific facet of internal mentation. Theoretically, the presence or absence of a self-directed thought could be easier to capture than internal processes related to the thought. From a participant standpoint, it may be more straightforward to introspectively identify shifts in attention than to self-assess one’s state of arousal in relation to arising thoughts.

### 6.4 Limitations and Future Directions

While we were able to validate the rt-fMRI-ES method through support for one of our hypotheses, future studies employing the rt-fMRI-ES method could benefit from several methodological refinements. One potential improvement could be to collect a higher number of trials per subject by either increasing the duration of each rt-fMRI-ES run or to include a larger number of runs in the study design. Our analyses using linear mixed-effects models were essentially a mean group estimate of the average effect of triggering trials from a region of interest for each subject. Thus, the overall effect of brain-state triggering was driven by the proportion of subjects that demonstrated the trend in their data. Collecting a higher number of trials per subject could provide more opportunity for a group-level desired effect to be observed.

Future implementations of the rt-fMRI-ES method could consider adopting alternative ways of operationalizing “high” or increased BOLD activity for triggering experience sampling. Our analysis of real-time brain activity involved calculating BOLD percent-signal-change for each volume relative to the region’s median BOLD signal at baseline. We defined BOLD activity as “increasing” if the BOLD signal displayed a consistent increase across three consecutive volumes. The number of volumes that our triggering algorithm used was decided based on simulations of our rt-fMRI-ES method using varying volume numbers (and assumed a TR of 2 seconds). Triggering based on BOLD increases across three volumes was found to present trials neither too liberally nor conservatively compared to four- or five-volume increases. Besides that rationale, the decision to trigger experience sampling from BOLD increase over a rigid number of volumes was relatively arbitrary.

There is no standard method for calculating ongoing real-time brain activity or for presenting stimuli from BOLD “activation” events. Prior studies have similarly used a 20-volume period to define baseline BOLD activity, but triggered stimuli based on a target area’s resting state activity being greater by at least one standard deviation than a control region’s activity (Hinds et al., 2013; Yoo et al., 2012). Another study presented trials of a task when ongoing mean BOLD activity across the past two volumes exceeded the 85^th^ percentile of the BOLD signal across a predefined sliding window (Chew et al., 2019). Our triggering algorithm was sufficient for detecting moments of *comparatively* decreased external-attention. However, using a percentile threshold based on a sliding window could potentially be more sensitive to capturing more dramatic fluctuations of brain activity, and consequently more extreme facets of ongoing thought (e.g. capturing low external-attention compared to relatively decreased external-attention).

### 6.5 Potential Applications and Conclusion

We validated the rt-fMRI-ES method for the purpose of capturing a specific form of spontaneous thought (decreased external-attention) through ongoing analysis of brain activity. This approach may pave the way for the prospective coupling of other aspects of ongoing thought to posited neural representations. While past groups have used EEG or physiological measures to trigger experience sampling (Hoemann et al., 2020; Siclari et al., 2017; Van Halem et al., 2020), the present study is the first to present thought probes based on ongoing BOLD activation during fMRI. Thus, our rt-fMRI-ES method offers improved spatial precision in the study of brain correlates of ongoing thoughts forms.

For future applications of rt-fMRI-ES, there are a few considerations to heed when selecting the target brain state to trigger experience sampling from. Selecting an empirically supported neural substrate of the thought form of interest, such as the PMC for internal mentation, has the potential to uniquely bolster existing evidence for a brain-thought relationship. On the other hand, selecting a more theoretically driven neural basis for the thought form of interest (e.g. as in our daIC analysis) could introduce a previously unexamined brain-thought relationship if support for the hypothesis is found. Additionally, future work could present experience sampling probes based on activity across multiple brain regions. For example, triggers could be derived from BOLD activity averaged over a functional network, or from using multi-voxel pattern analysis distributed over one or more regions. These approaches may be more robust than triggering experience sampling from activity constrained to a single region of interest, as specific facets of spontaneous thought are likely to originate from activity distributed across multiple brain areas (Christoff et al., 2016; Kim et al., 2024; Kucyi et al., 2023). Overall, decisions regarding the operationalization of a “brain state” are highly specific to the analytical aims of the study and to the form of thought that rt-fMRI-ES is being used to capture.

The rt-fMRI-ES method can link brain states to various types of ongoing thought in many ways. Future work could use rt-fMRI-ES to index ongoing engagement in thought or the dynamic movement of thought, enabling the potential to better understand neural signatures of both automatic and deliberate spontaneous thought constraints. Presenting probes relevant to thought continuity or repetition could elucidate the neural bases of ongoing thought progressions, or what occurs in the brain when one thought ends and another begins. Using rt-fMRI-ES to sample clinically-relevant forms of thought, such as perseverative thought or obsessive thought, could facilitate the identification of specific brain states as potential biomarkers for cognitive patterns pertinent to psychiatric disorders. In the future, the rt-fMRI-ES method could be used to identify targets for neuromodulation therapies for a variety of mental disorders.

Spontaneous thoughts are a ubiquitous part of the human experience. Studying the brain mechanisms underlying these thoughts allows us to understand how our brains self-generate mental content in the absence of external stimuli. Using rt-fMRI-ES to track mental experiences as they unfold may provide unique insights into the neural bases of unconstrained thought forms.

## Author Contributions (CRediT Statement)

**Tiara Bounyarith:** Writing – original draft, Formal analysis, Project administration, Data curation, Visualization, Writing – review and editing, Investigation

**David Braun:** Writing – original draft, Formal analysis, Visualization, Writing – review and editing

**Aaron Kucyi:** Writing – original draft, Conceptualization, Methodology, Supervision, Funding acquisition, Software, Writing – review and editing, Investigation

## Acknowledgements

This work was supported by the National Institute of Mental Health of the National Institutes of Health under award number R21MH129630 to Aaron Kucyi and a Graduate Research Fellowship from the National Science Foundation awarded to Tiara Bounyarith. The content is solely the responsibility of the authors and does not necessarily represent the official views of the National Institutes of Health.

## Data and Code Availability

In accordance with requirements set by the project sponsor, all study raw data will be available through the online National Institutes of Mental Health Data Archive (NDA), DOI: https://dx.doi.org/10.15154/5aka-y257.

Code and data used for analysis and visualization are available on OSF, https://osf.io/as2q9/overview?view_only=cb859cc4666842c7b347f9802d535d6c.

## PCI-RR Study Design Chart

| Question | Hypothesis | Sampling Plan | Analysis Plan | Rationale for deciding the sensitivity of the test for confirming or disconfirming the hypothesis | Interpretation given different outcomes | Theory that could be shown wrong by the outcomes | Observed Outcome |
| --- | --- | --- | --- | --- | --- | --- | --- |
| Does spontaneous BOLD activation in the daIC predict the occurrence of mental experiences that are characterized by high subjective arousal ? | H1: We predict that subjective ratings of arousal time-locked to daIC-activation trials will be higher than ratings time-locked to non-daIC-activation trials (including both PMC-activation trials and time-out trials). This hypothesis is rooted in evidence for a role of the daIC in assigning salience to spontaneous thoughts and in physiological arousal. | Power analysis was conducted using a previous study we conducted that used fMRI and random-interval experience sampling. Based on power simulations using the effect size we identified in that study, an N of 53 is required for .90 power. Given rt-fMRI-ES is intended to boost statistical power relative to random-interval experience sampling protocols, we consider this to be a conservative | A linear mixed-effects model will be conducted with trial-types (daIC activation-triggered, PMC activation-triggered, or “time-out” non-brain-triggered) set as fixed effects, with random effects grouped by participant, and self-reported experience sampling arousal ratings as the outcome variable. | Based on prior fMRI literature inferring associations between spontaneous brain activity and experience sampling reports, setting the significance level at $P < 0.05$ will be sufficient to detect significant contrasts between the levels of our main predictor. | If daIC triggered trials are associated with significantly higher self-reported arousal ratings relative to the other two trial types, then our results will validate our rt-fMRI-ES technique by demonstrating its ability to capture salient spontaneous experiences using real-time BOLD analysis. This finding would further corroborate prior evidence that the daIC is linked to subjective arousal.<br>If daIC activation triggered trials are not associated with significantly higher self-reported arousal ratings relative | If our results support our hypothesis regarding daIC activation levels predicting high-arousal mind-wandering experiences, this finding would bolster insight into the role of non-default mode network brain regions in self-generated experiences.<br>If our results do not support our hypothesis regarding daIC activation levels predicting high-arousal mind-wandering experiences, this finding would encourage | We did not find evidence for our hypothesis that arousal ratings would be higher for daIC-triggered trials compared to non-daIC-triggered trials. Across 1243 trials from 42 participants eligible for this linear mixed-effects model, there was no statistically significant difference between predicted mean arousal time-locked to daIC-triggered trial compared to non-daIC-triggered trials ($b = -0.02$, 95% CI $[-0.25, 0.20]$, $df = 39.7$, $t = -0.21$, $p = .84$).<br>We interpret this finding as potential evidence for a difference between subjective self-reported arousal and task-evoked physiological arousal – an arousal construct commonly associated with daIC involvement. |
|  |  | target sample size, and we will aim to collect data from 60 participants to account for participants who fail to meet our criteria for inclusion. |  |  | to the other two trial types, we will interpret this as a negative finding. These negative findings would constrain hypotheses to guide future work into refining real-time BOLD analysis, and studying the role of the daIC in subjective arousal. | further rigorous study into the salience network's role in self-generated experiences. Such a finding could also motivate either the development of a more efficient approach of real-time BOLD analysis of the daIC, or the study of a different salience network node in association with mind-wandering. |  |
| Does spontaneous BOLD activation in the PMC predict the occurrence of internally-focused mind-wandering episodes? | H2: We predict that subjective ratings of attention to inner mental experience time-locked to PMC-activation trials will be higher than ratings time-locked to non-PMC-activation trials (including both daIC-activation trials and | Same as H1. | A linear mixed-effects model will be conducted with trial-types (daIC activation-triggered, PMC activation-triggered, or "time-out" non-brain-triggered) set as fixed effects, with random effects grouped by | Same as H1. | If PMC activation triggered trials are associated with significantly higher self-reported internal attention ratings, then our results will validate our rt-fMRI-ES technique by demonstrating its ability to capture the onset of self-referential spontaneous experiences using real-time | If our results support our hypothesis regarding PMC activation levels predicting increased internally-focused attention, this finding would build upon previous work connecting DMN regions to mind-wandering via retrospective data analyses, | We found support for our hypothesis that self-reported internal attention would be higher (i.e. external-attention would be lower) for trials time-locked to PMC activation compared to other trial types. Across 1429 trials from 49 participants eligible for this model, external-attention ratings were significantly lower for trials time-locked to PMC activation compared to other trial types ($b = -0.30$, 95% CI $[-0.55, -0.06]$). |
|  | time-out trials). This is supported by a theorized role of the PMC in stimulus-independent thought, regardless of specific content. |  | participant, and self-reported experience sampling attention ratings as the outcome variable. |  | BOLD analysis. This finding would further corroborate prior evidence that the PMC is linked to internally-focused cognition.<br>If PMC activation triggered trials are not associated with significantly higher self-reported internal attention ratings, we will interpret this as a negative finding. These negative findings would constrain hypotheses to guide future work into refining real-time BOLD analysis, and studying the role of the PMC and other DMN nodes in internally-focused attention | offering plausible real-time fMRI evidence that mind-wandering episodes can be prospectively predicted from ongoing brain activity.<br>If our results do not support our hypothesis regarding PMC activation levels predicting increased internally-focused mind-wandering experiences, this finding would encourage further rigorous study into the PMC's role in self-generated experiences. Such a finding could motivate either the development of a more efficient approach of real-time BOLD analysis of the PMC, or the study of a<br>different<br>DMN node in<br>association<br>with mind-wandering. | $df = 45, t = -2.50, p = .016$).<br>We interpret this finding as a validation of our rt-fMRI-ES method's ability to capture the onset of internally-oriented thoughts using real-time analysis of ongoing BOLD activation.<br>Furthermore, we consider this finding to bolster the classically-found relationship between internally-oriented thought and increased PMC activation. |

